# Development of a Scoring Function for Comparing Simulated and Experimental Tumor Spheroids

**DOI:** 10.1101/2022.08.08.503266

**Authors:** Julian Herold, Eric Behle, Jakob Rosenbauer, Jacopo Ferruzzi, Alexander Schug

**Author notes:** These authors contributed equally to this work (Julian Herold, Eric Behle).

## Abstract

Enormous progress continues in the field of cancer biology, yet much remains to be unveiled regarding the mechanisms of cancer invasion. In particular, complex biophysical mechanisms enable a tumor to remodel the surrounding extracellular matrix (ECM), thus allowing cells to escape and invade alone or as multicellular collectives. Tumor spheroids cultured in collagen represent a simplified, reproducible 3D model system, which is sufficiently complex to recapitulate the evolving internal organization of cells and external interaction with the ECM that occur during invasion. Recent experimental approaches enable high resolution imaging and quantification of the internal structure of invading tumor spheroids. Concurrently, computational modeling enables simulations of complex multicellular aggregates based on first principles. The comparison between real and simulated spheroids represents a way to fully exploit both data sources, but remains a challenging task. We hypothesize that comparing any two spheroids requires first the extraction of basic features from the raw data, and second the definition of key metrics to match such features. Here, we present a novel data-agnostic method to compare spatial features of spheroids in 3D. To do so, we define and extract features from spheroid point cloud data, which we simulated using Cells in Silico (CiS), a high-performance framework for large-scale tissue modeling previously developed by our group. We then define metrics to compare features between individual spheroids, and combine all metrics into an overall deviation score. Finally, we use our features to compare experimental data on invading spheroids in increasing collagen densities. We propose that our approach represents the basis for defining improved metrics to compare large 3D data sets. Moving forward, this approach will enable informing *in silico* spheroids based on their *in vitro* counterparts, and vice versa, thus enabling both basic and applied researchers to close the loop between modeling and experiments in cancer research.

**Author summary:** Cells within a tumor use various methods to escape and thereby invade into healthy parts of the body. These methods are studied experimentally by examining tumor spheroids, spherical aggregates of hundreds to thousands of individual cells. Such spheroids can also be simulated, and the comparison of simulation and experiment is desirable. Here, we present an analysis strategy for the comparison of tumor spheroids, a widely used workhorse of cancer research. Using this strategy, we aim to improve the collaborative potential between experimentalists and theorists.

## 1 Introduction

The worldwide challenge of fighting cancer is as urgent as ever [1, 2]. When trying to understand the mechanisms driving the disease, one is faced with a complex and wildly inhomogeneous landscape of cellular properties and interactions, which vary both within and between cancer types [3–5]. Furthermore, cancer is not one single disease, but rather refers to a large number of diseases with shared characteristics, those characteristics are captured in the hallmarks of cancer [2, 6]. The processes underlying these diseases, such as the rise of malignancy via loss of adhesion and subsequent increased motility [7], span a wide range of scales, both in space and in time [8]. To further the understanding of cancer, it is crucial to decipher how these processes interact and lead to the formation of macroscopic invasive tumors. Thus, combating cancer requires input from many different domains of science, such as biology, medicine, and pharmacology, but also physics, computer science, and mathematics [8, 9]. Unfortunately, time-resolved analysis of *in vivo* tumor tissue is challenging, as due to low spatial or temporal resolution of imaging methods, single-cell resolution 4D trajectories are not yet widely applicable. To increase accessibility for analysis, the system has to be divided into smaller subsystems. Thus, *in vitro* and *in silico* models are created, allowing the study of individual aspects of the system. An *in vitro* example is the study of tumor spheroids, which represent a useful model system for studying tumor growth and cell dynamics [10]. Tumor spheroids are spherical arrangements of hundreds to thousands of cells, which can be placed within a structural extracellular matrix (ECM), e.g. a collagen scaffold. They are widely used for studying e.g. drug response, tissue fluidity and tumor invasion [11–13]. On the *in silico* side, tumor growth models of varying degree of coarse-graining are being developed [14–16], some of which are also applied to simulate tumor spheroids [17, 18]. Thus, both experimentalists and theorists generate data for the same systems, but these studies are usually not compared quantitatively. Quantitative comparison is an important step towards fully leveraging the results of both groups and requires an adaptive and robust comparison strategy for spheroid data, regardless of its origin. To our knowledge however, there is currently no data-agnostic strategy for systematically comparing two spheroids (see also figure 1). Hence, in this study, we want to provide a toolbox of features which may be extracted from a given 3D structure of a spheroid, and metrics to compare these extracted features between different spheroids. These features and metrics can be used on their own, or in combination, to obtain an overall deviation score. Our strategy utilizes point cloud data, in which each point denotes the position of a cell. Importantly, this enables the comparison of both simulated and experimental spheroids. To demonstrate this, we applied our toolbox to previously published data that captured structural differences in triple-negative breast cancer spheroids invading into a collagenous ECM of varying density [13] (see Fig 2 a). Such experimental data were used as a motivation to simulate a variety of spheroid behaviors in silico (see Fig 2 b). Simulations were performed using our previously developed platform “Cells in Silico” (CiS) [19]. CiS is a highly scalable general-purpose framework for tissue simulation at subcellular resolution. It extends a cellular Potts model with an agent-based layer, and allows the description of various properties such as cell-cell adhesion, cell compressibility and cell motility, and phenomena such as cell divison, cell mutation and ECM interactions, among others (see also Section 4.1). We performed a large set of simulations of 3D tumor spheroids in ECM, spanning a wide range of parameters (see Section 4.2 for details). For this study, we want to highlight a representative subset of four spheroid phenotypes, that we found in the simulated data: “Spherical”, “Spherical with far gaslikes”, “Deformed”, and “Disordered”. These phenotypes emerged from various combinations of parameters, including ECM density, ECM degradation, cell motility, and cell-cell adhesion (see Section 4.3). They will be used as examples throughout.

**Figure 1:**
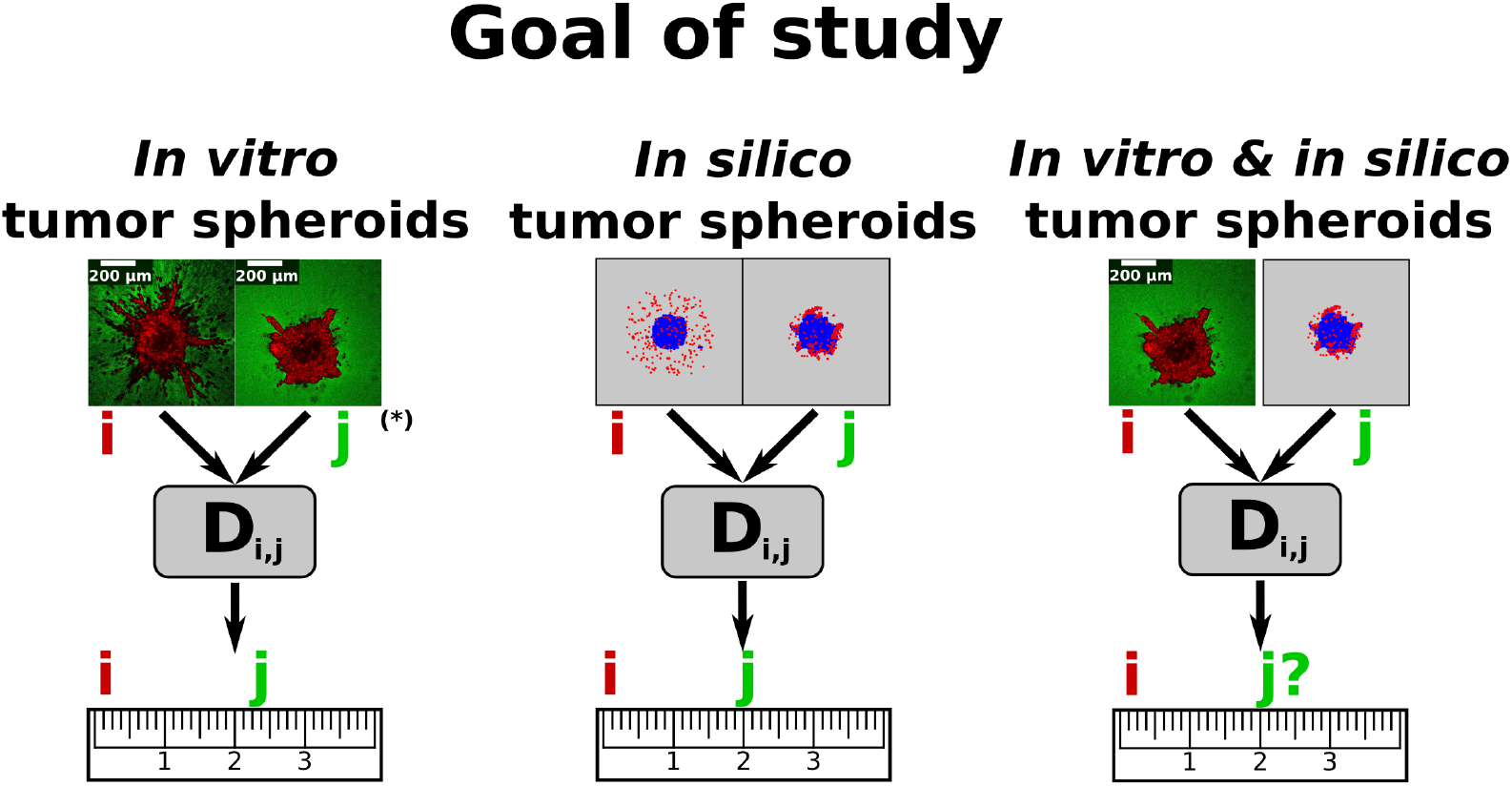
Scientific context. Tumor spheroids are one of the main workhorses of tumor tissue analysis. Both experimentalists and theorists generate data from such spheroids, but to our knowledge there is currently no unified comparison strategy. This study deals with defining data features and comparison metrics, which can be combined into an overall deviation score *D*_i,j_ between individual spheroids *i* and *j*. (*) Spheroid images provided by Kang et al. [13].

**Figure 2:**
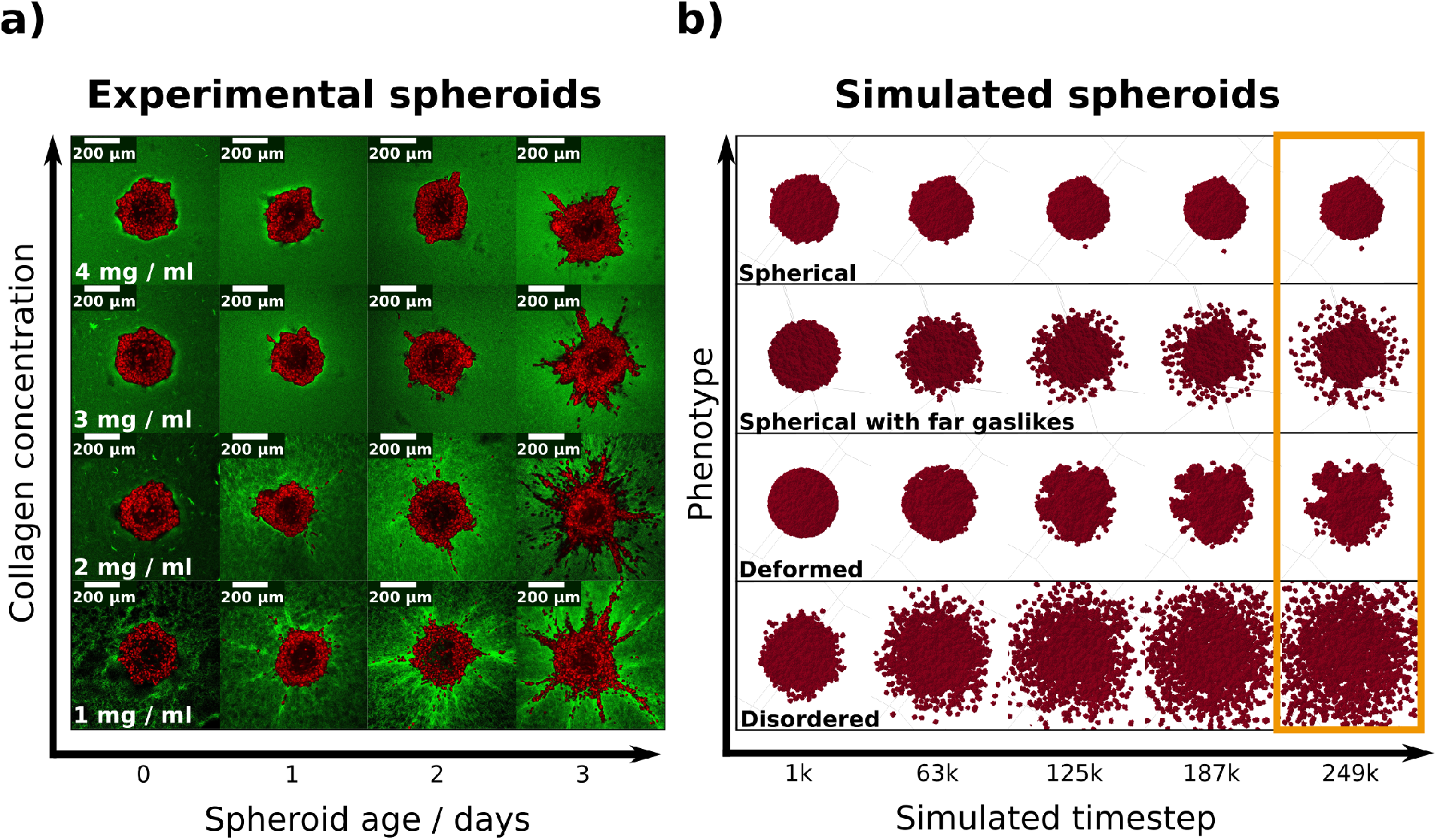
**a)** 2D cross-sections of 3D multiphoton microscopy image stacks depicting the spatio-temporal evolution of MDA-MB-231 spheroids in increasing collagen concentrations [13]. Spheroids were imaged at zero, one, two or three days after embedding in collagen, and were then fixed and imaged (see Section 4.4). **b)** Time evolution of simulated spheroids displaying four different phenotypes (the order is unrelated to a)): : “spherical”, “spherical with far gaslikes”, “deformed”, “disordered”. Each phenotype resulted from different combinations of parameters connected to the cell motility, the cell-cell adhesion, the interaction with the ECM, and the ECM density (see Section 4.3). Each simulation lasted 250 000 Monte-Carlo (MC) steps, and shown are five snapshots for each simulation. Throughout this study, we focused on the final configuration (orange rectangle), and used replicates from each phenotype.

In the following, we will first describe the spatial features that we extracted from individual experimental and simulated spheroids. Next, we will discuss our strategy for comparing these features between multiple spheroids, including the derivation of an overall deviation score, and how it can be tuned for a specific use-case. After validating our strategy via a transformation study, we will show comparisons between exclusively simulated spheroids, exclusively experimental spheroids, and comparisons between simulated and experimental spheroids. We will conclude by evaluating the success of our method, and providing an outlook for its further use.

## 2 Results

### 2.1 Individual features

Our study focused on the analysis of spatial properties of tumor spheroids. We utilized data containing the three-dimensional positions of all individual cell centers at one given point in time. For the analysis of these data we considered various features which could be used individually or as an overall deviation score (see Section 2.3). These features and their applicability are highlighted in the following.

#### Cell density distribution

Analyzing the distribution of cell density is useful for determining the extent of the bulk of the spheroid, as well as its geometry. For our studies we focused on the so-called central local density, which we defined as the fraction of cells found within spherical layers of constant thickness and increasing radii around the spheroid center. For a uniform spherical distribution of cells, this density is non-zero only within the spheroid bulk (see Fig 3 a). Disordered spheroids, on the other hand, exhibit a distribution over a larger domain.

**Figure 3:**
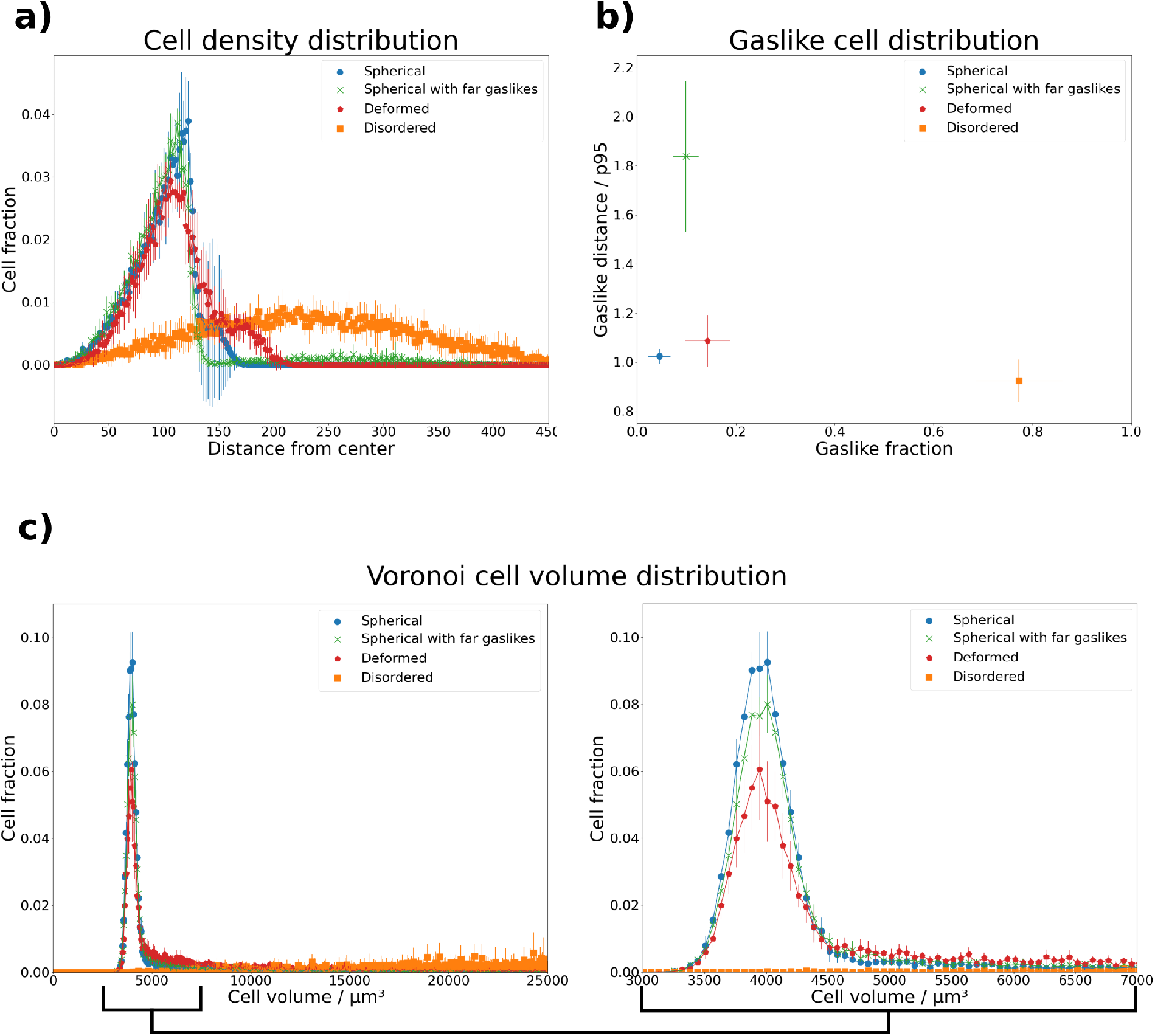
Visualization of cell based features. **a) Cell density distribution**. Shown is, for each phenotype, the average fraction of cells within spherical layers around the spheroid center *versus* the radii of these layers. The “spherical” phenotype shows a steep drop, while the “deformed spheres” and “spheres with gaslikes” phenotypes show a long-tailed distribution. **b) Gaslike cell distribution**. Shown are the average fractions of gaslike cells according to equation 1 *versus* their normalized average distance to the spheroid center. The fraction of gaslikes exhibited by the “spherical”, “spherical with far gaslikes” and “deformed” phenotypes is similar, but the distance from the spheroid center is far greater for the “sphericals with far gaslikes” phenotype. The “disordered” phenotype on the other hand contains many cells classified as gaslikes across the entire spheroid volume. Their normalized average distance from the center evens out to a value slightly below 1. **c) Voronoi cell volume distribution**. Shown are histograms of the average Voronoi cell volumes found in the four phenotypes.The “spherical” phenotype shows a sharp peak around a volume of 4000 µm^3^. The “spherical with far gaslikes” and “deformed” phenotypes show a similar behavior, with a more pronounced tail towards larger volumes. Finally, the “disordered” phenotype shows volumes distributed over a wide range. Values between 7500 µm^3^ and 20000 µm^3^ were cut in order to highlight the differences between phenotypes in the peak around 4000 µm^3^.

#### Gaslike cell distribution

The detachment of single “gaslike” cells from a spheroid has been a recent focus [13] and can be used to distinguish between ordered and disordered spheroids. However, the assignment strategy of the “gaslike” status needs to be well defined. Kang et al. were able to experimentally measure the spheroid boundary [13], and defined cells outside of this boundary as “gaslike”. Since the spheroid boundary is not tracked in our simulations, we used a definition based on nearest-neighbor distances and distance from the center of the spheroid. The set of gaslike cells *G* as a subset of all cells *C* is thus defined as follows:

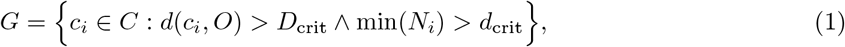

where *D*_crit_ is the threshold distance from the spheroid center *O, N*_*i*_ = {*d*(*c*_*i*_, *c*_*j*_) | *c*_*j*_ ∈ *C, c*_*j*_ ≠ *c*_*i*_} is the set of Euclidean distances between cell *c*_*i*_ and all other cells, and *d*_crit_ is the threshold neighbor distance. The first constraint in equation 1 provides the context of a bulk structure, and its parameter *D*_crit_ can be selected considering the inflection point of the central local density. The second constraint ensures that only detached cells are defined as gaslikes, and its parameter *d*_crit_ was chosen by considering the mean distance between all cells. Fur our purposes we selected the data by Kang et al. [13] as basis for deriving these parameters: *D*_crit_ = 125 µm, *d*_crit_ = 16 µm. This set *G* can be used to compute various properties, such as the fraction of gaslikes and their average distance from the spheroid center. We combined these two properties in this feature, defining it as a point *p* within the space spanned by them. The first property *p*_x_ describes the fraction of all cells in the spheroid that are detached. The second property *p*_y_ describes the mean distance of the detached cells from the spheroid center.

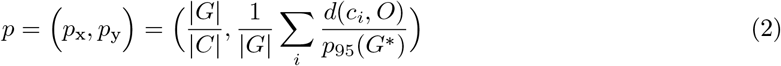

where *p*_95_(*G*^*^) is the 95th percentile of distances of the non-gaslike cells *G*^*^ = *C \ G* from the spheroid center, which serves as a normalization factor. We included only the non-gaslike cells for this normalization factor, because our aim was to define the distance relative to the spheroid bulk. As shown in Fig 3 b, the “disordered” and “spherical with far gaslikes” phenotypes can be distinguished from the others using this feature, but it is suited less well for comparing “spherical” and “deformed” spheroids.

#### Voronoi cell volume distribution

The distribution of Voronoi cell volumes within the spheroid serves as a measure of cell deformation, as well as their confinement. To obtain these volumes, we performed a Voronoi tessellation [20] on the cell center point cloud, during which the system was divided into *N*_cells_ regions according to the distances between adjacent cells. It is important to note, that the voronoi cell volumes are not the same as the biological cell volumes, but represent a proxy where detached cells occupy a significantly larger volume. We generated a histogram of the voronoi cell volumes, as shown in Fig 3 c). Here we observe that the first three phenotypes are distributed sharply around a volume of roughly 4000 µm^3^ with a tail, while the volumes of the “disordered” phenotype are evenly distributed over a much wider range. The tail of the first three distributions is a useful artefact of the Voronoi tesselation, as it allows to extract additional information about the bulk spheroid surface.

#### Spheroid surface and surface deformation

While the cell density distribution provides a measure of the spheroid size, its information about the spheroid shape is limited. To study this in more detail, we need to approximate the spheroid surface, as we want to distinguish deformed spheroid bulk from spherical bulk. We did this via surface triangulation using the marching cubes algorithm [21]. To apply this algorithm we performed some preprocessing of the point cloud data: first, we extracted the set of non-gaslikes *G*^*^, as we were only interested in the shape of the spheroid bulk. Next, we voxelized our data to obtain a continuous spheroid volume. From this, the surface could be triangulated via the marching-cubes algorithm. Finally, we extracted two features from this: first, we calculated the surface area from the triangle mesh (see Fig 4 a)), and second, we analyzed the surface deformation by investigating the orientations of the mesh vertices. This was done by calculating, for each vertex, the scalar product between its normal vector and its origin vector, with the spheroid center at the origin. Then these scalar products were combined in a histogram. The vertex orientations serve as a measure of deformation, since for a perfect sphere all scalar products are equal to 1, and a deformed sphere results in a more widely spread distribution (see Fig 4 b).

**Figure 4:**
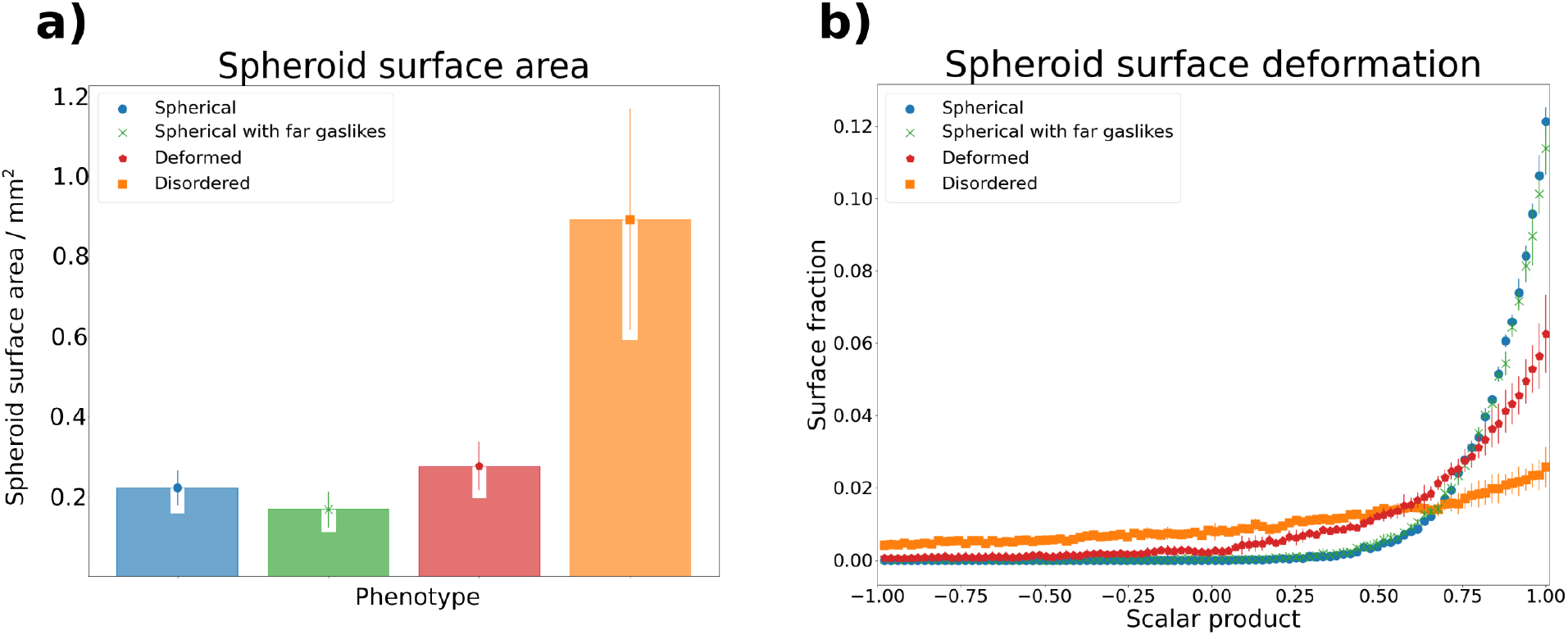
Visualization of spheroid bulk based features extracted from simulation data for four different phenotypes. Surface information was extracted via the marching cubes algorithm [21]. **a) Spheroid surface area**. Shown is the average surface area found for each phenotype. The “spherical with far gaslikes” phenotype has the smallest average surface area, due to the spherical bulk containing less cells than that of the “spherical” phenotype. The larger average surfaces of the other two phenotypes are due to the more irregular shape of the bulk. **b) Spheroid surface deformation**. Shown are histograms of the scalar products between vertex normal vectors and vertex origin vectors, with the origin denoting the center of the spheroid. The vertices were obtained from surface triangulation of the spheroid point cloud. The two spherical phenotypes exhibit a sharp peak at scalar products of 1, while the deformed phenotypes are spread more widely.

By using these features we were able to measure and quantitatively describe different aspects of individual tumor spheroids. This provided a basis on which we could compare two spheroids with each other. Such a comparison required the definition of distance metrics for each feature, which are highlighted in the following.

### 2.2 Individual metrics

To accomodate the different types of output data between features, we required suitable metrics. For the spheroid surface area, which provided a scalar value per spheroid, we used the mean squared error (MSE). For the gaslike cell distribution, which provided a tuple of two coordinates per spheroid, we used the Euclidean distance. Finally, for the distribution-based features, i.e. cell density distribution, Voronoi cell volume distribution and spheroid surface deformation, we used the 1-Wasserstein distance (WSD). The Wasserstein distance is a metric between probability distributions, and is a common sight in mathematics, especially statistics and computer science. The *p*-Wasserstein distance between two probability measures *µ* and *ν* on ℝ^*d*^ is defined as:

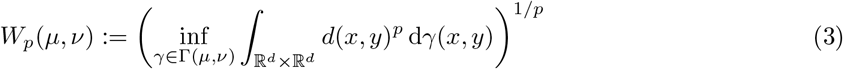

where Γ denotes the collection of all joint probability measures *γ* with marginals *µ* and *ν* [22].

An intuitive illustration of the Wasserstein distance can be given by viewing each distribution as a pile of earth of different shape, and considering the amount of work that has to be done to transform one pile into the other. Assuming this work to be equal to the product between the amount of earth that has to be moved and the distance it needs to be moved, the Wasserstein distance is the minimum amount of work that has to be done. Due to this illustration, the WSD is often referred to as “earth mover’s distance” [23].

### 2.3 Combination of multiple metrics

At this point, the individual features described in the previous sections could be reliably compared between two spheroids using our defined metrics. Next, one of our main goals was to combine these features and metrics into a single scalar value, which could then serve as an overall deviation score between two spheroids. Many different questions regarding the comparison of tumor spheroids require such a singular scalar. From an experimentalist’s view, this could be the comparison of spheroids cultivated in different conditions, with the goal of quantitatively determining how the change of one experimental variable influences the spheroid growth and invasion pattern. A problem faced by theorists running simulations is how to optimize the model parameters to reproduce experimental results. Both problems require one scalar distance measure like the one we aimed to derive here. Before doing so, we need to address the fact that it is unlikely for such a distance measure to be generally applicable to all types of tumor spheroids and experimental settings. This is due to the high dimensionality of even a single spheroid dataset at a single point in time. To illustrate this, we consider the case of comparing two spheroids, each containing 1000 cells. The desired distance is a function *f* : ℝ^3000^ × ℝ^3000^ → ℝ. Such a function will, by design, project many different pairs of spheroids onto the same point in ℝ. This property can hardly be circumvented, and is desired in a distance measure. On the other hand, this also means that the measure has to be carefully selected depending on the use case. Therefore, in addition to defining the overall deviation score here, we will also propose a method to adapt the score to different use cases.

#### Standardization

We first ensured that all metrics were on a similar scale. For this standardization, we used five replicates from each of the four phenotypes from our simulations, and compared each feature, resulting in *N*_spheroids_ = 20 spheroids and *N*_distances_ = 400 metric distances per feature *f*. These distances *d*_i,j,f_ between spheroid *i* and spheroid *j* were then transformed according to equation 4:

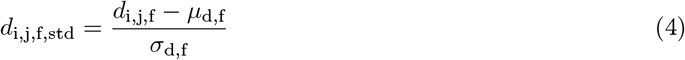

where *µ*_*d,f*_ and *σ*_*d,f*_ respectively denote the mean and standard deviation across the *N*_distances_ values for each feature *f*. Since this standardization may lead to values of *d*_i,j,f,std_ below zero, and we aimed to define a positive distance for each feature, we further shifted each value by the minimum across all *d*_i,j,f,std_, finally arriving at 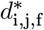 as defined in equation 5:

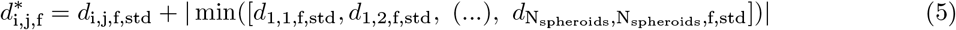

#### Overall deviation score and use case adaptation

Next, we defined the overall deviation score *D*_i,j_. This definition entailed merging the previously standardized 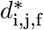 via the following linear combination:

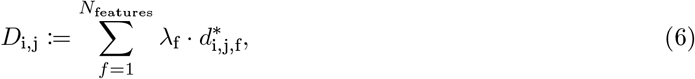

where *λ*_f_ denote the weight factors for each feature, i.e. how much it contributes to the final deviation score *D*_i,j_. In order to optimize these values, we once again turned to our simulated phenotypes and their five respective replicates. We required values of *λ*_f_ that minimized the intra-phenotype deviations and maximized the inter-phenotype deviations. This can be formulated as a maximization problem for the following relation:

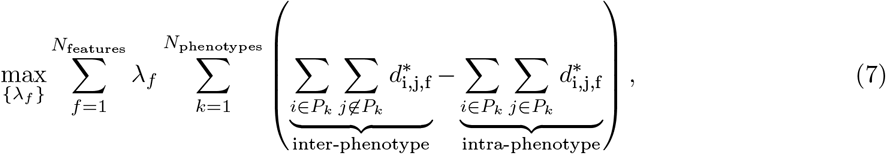

where *P*_*k*_ is the set of all spheroids of phenotype *k*. This optimization procedure can be interpreted as an inverse clustering. During clustering, the property described in equation 7 is maximized by assigning individuals to a cluster. In contrast to this, our method uses prior clustering information to optimize the metric space itself. This shows resemblance to methods of contrastive learning, with the notable difference that we use a linear model in our approach [24]. Assuming that phenotypes are correctly grouped, maximizing Equation 7 already ensures that each *λ*_*f*_ *>* 0. Additionally, we decided on the following constraint:

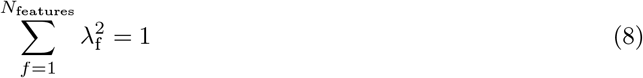

This constraint is important to prevent the optimization procedure from collapsing towards the trivial solution of setting *λ*_*f*_ → ∞. It also fixes each *λ*_*f*_ to the domain [0, 1]. Furthermore, using the square of each *λ*_f_ minimizes the influence of outliers across features. The contribution of each feature to the overall deviation score resulting from the optimization of the *λ*_f_ is shown in Table 1.

**Table 1:**
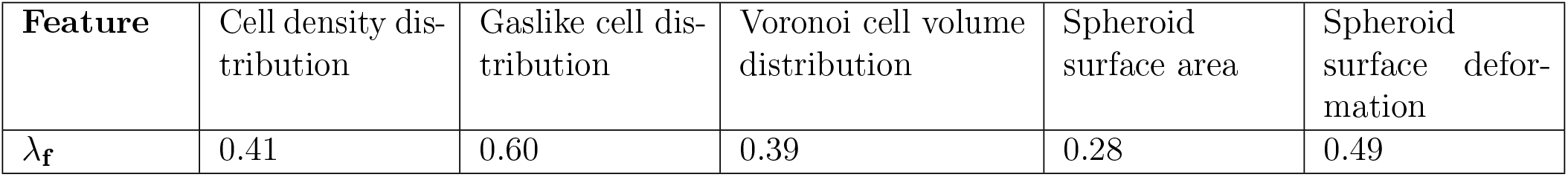
Fitted weight factors for each feature contributing to the overall deviation score between two spheroids (see equation 6). The values were obtained by maximizing equation 7, with the simulated phenotypes serving as a calibration set (see Section 4.3).

### 2.4 Validation: transformation study

Validating a metric, such as the one derived in this work, requires a set of data examples with known relation. For this purpose, we designed a transformation study, in which we generated multiple point clouds from a reference spheroid using transformation functions. These functions were selected in such a way, that for a useful metric we expected higher distances between reference and transformed point cloud for higher transformation strengths. On the other hand, the metric has to remain invariant under transformations related to the frame of reference, e.g. rotation or translation of the point cloud. Therefore, we also included these transformations. Hence, we tested for the following properties:

1. invariance under rotation and translation,
2. monotony within the domain of interest: a small deviation from an original spheroid shall result in a lower distance than a large deviation.

We investigated these properties for the five described features and the overall deviation score by appling four transformations to a spheroid 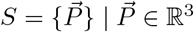 of the “spherical” phenotype. The transformations were represented by functions in the space of point clouds *T* : *S* × *α* → *S*. With this approach, we aimed to verify both of the above properties. Invariance is shown, when the distance does not depend on the strength of the transformation. Similarly, monotony is shown, when the distance metric grows monotonously with the transformation strength. The four transformations that we used were the following:

#### Rotation

Rotating each cell of the spheroid by a given angle *α* around an arbitrary axis *i*:

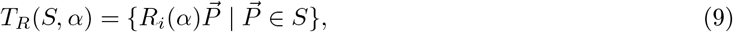

where *R*_*i*_(*α*) is the rotational matrix.

#### Noise

Adding a random vector drawn from a normal distribution to the position of each cell:

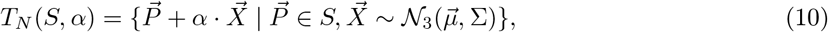

#### Deformation

Translating each cell along the radial vector of the spheroid, modulated by the spherical angles of the cell’s position:

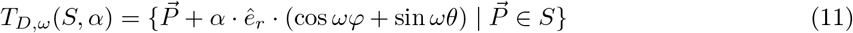

This deformation can be interpreted as adding ripples with frequency *ω* and amplitude *α* to the spheroid surface.

#### Scaling

Multiplying the position of each cell by its distance to the center of the spheroid:

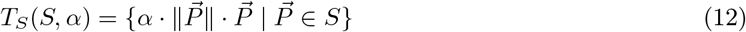

This transformation affects cells with a larger distance to the spheroid center more strongly than those close to the it. The spheroid density is therefore not conserved.

A visual example for each transformation is provided in Fig 5. Translational invariance was ensured, since all features depend only on relative and not absolute distances. Rotational invariance was expected due to the rotational symmetry of the underlying features. Nonetheless, we wanted to test whether artifacts, produced by the voxelization for features related to the spheroid surface (see Section 2.1), had any notable effect. For this reason, we included the rotation transformation. The remaining transformations were chosen to validate the monotony of the deviation score.

**Figure 5:**
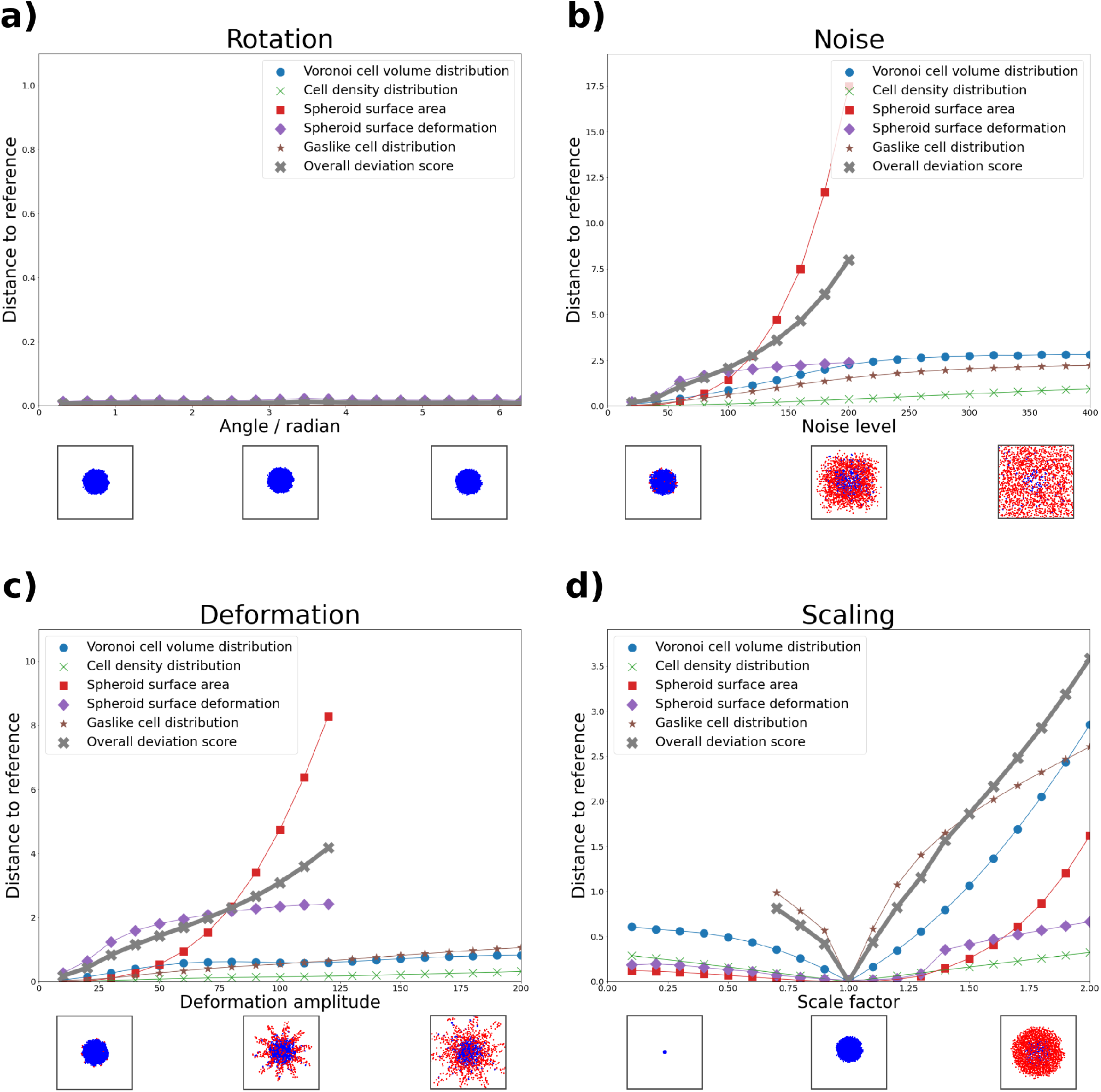
Feature comparison for spheroid point clouds resulting from four different transformation functions. Shown are the standardized metric distances between the un-transformed reference spheroid and an increasingly transformed version for each data feature. In addition, the combined deviation score is depicted in gray crosses for each transformation (see Section 2.3). Below each subfigure, we provide a top-down view snapshot of the spheroid at three levels of transformation. Blue cells are classified as non-gaslike, and red cells are classified as gaslike. **a) Rotation**. Except for negligible changes in the spheroid surface deformation feature, we observe no change at increasing rotation angle. This supports rotational invariance of our features. **b) Noise**. For each feature, the distance increases at increasing noise level, most rapidly for the spheroid surface area. Due to the loss of a solid core at high noise levels, the spheroid surface area and deformation features are no longer sensible, and were therefore cut. The deviation score increases approximately linearly up to a noise level of 120, at which point the quadratically increasing spheroid surface area becomes dominant. **c) Deformation**. Similar behavior to b) is observed here. Above a deformation amplitude of 100 the spheroid again loses its solid core, and surface values were therefore cut. The deviation score increases approximately linearly up to a deformation amplitude of 120. **d) Scaling**. We observe increased distances both for scale factors below and above 1. Due to the fixed values of *D*_crit_ and *d*_crit_ (see equation 1), the gaslike distribution feature is scale-dependent, and strongly varies here. For scale factors below 0.6, no gaslikes were found, and therefore the values of this feature were cut. The deviation score increases approximately linearly both for scale factors smaller and larger than 1.

We applied each transformation at increasing strength and compared the resulting spheroid with the untransformed version. The results of this are shown in Fig 5. Starting with the rotation transformation in subfigure a), we observed no change in the feature distances at increasing rotation angle, except for negligible changes in the spheroid surface derformation. This underlines the rotational invariance of our features. For the other three transformations, we observed monotony in all cases. Those features related to the spheroid surface could not be meaningfully extracted when the spheroid bulk was disrupted, i.e. at high degrees of the noise and deformation transformations (subfigures b) and c)). Similarly, the gaslike distribution feature was undefined for small values of the scaling transformation (subfigure d)), since no cells were classified as gaslike here. Aside from these edge-cases, our features behaved robustly. It is interesting to note that the overall deviation score scaled approximately linearly with the transformation strength, excluding the aforementioned extreme cases. This property can be viewed as a stronger version of the monotony property. Importantly, this was not used as a constraint when optimizing the weights, but emerged from the procedure itself.

### 2.5 Validation: comparing simulated spheroid phenotypes

As a second way to validate our methods, we now moved to the comparison of simulated spheroids. We chose the final simulation state, after 250 000 MC steps, of five new replicates from each phenotype. Importantly, these were not the same replicates which we used earlier for the calibration of the weight factors. We calculated the overall deviation score for each pair. As shown in Fig 6 a), we compared individual replicates (upper triangle), and we also combined replicate comparisons into an average phenotype deviation score (lower triangle). Importantly, the “disordered” phenotype is not a biologically occuring configuration, but occasionally arose during our exploration of the simulation parameter space. It was thus crucial for us to include this phenotype explicitly. Since it deviates strongly from the others, we have stretched the color bar for deviation score values below 1.5 for illustrative purposes. The deviation score was lowest when a phenotype was compared with itself, and highest, when any phenotype was compared with the “disordered” one. The “spherical”, “spherical with far gaslikes” and “deformed” phenotypes showed a smaller deviation score between each other, but were nonetheless distinguishable. This is underlined in subfigure b), in which we show box plots of the overall deviation scores between the “spherical” phenotype and the others. Here, each phenotype comparison was clearly distinct from the comparison of the “spherical” phenotype with itself.

**Figure 6:**
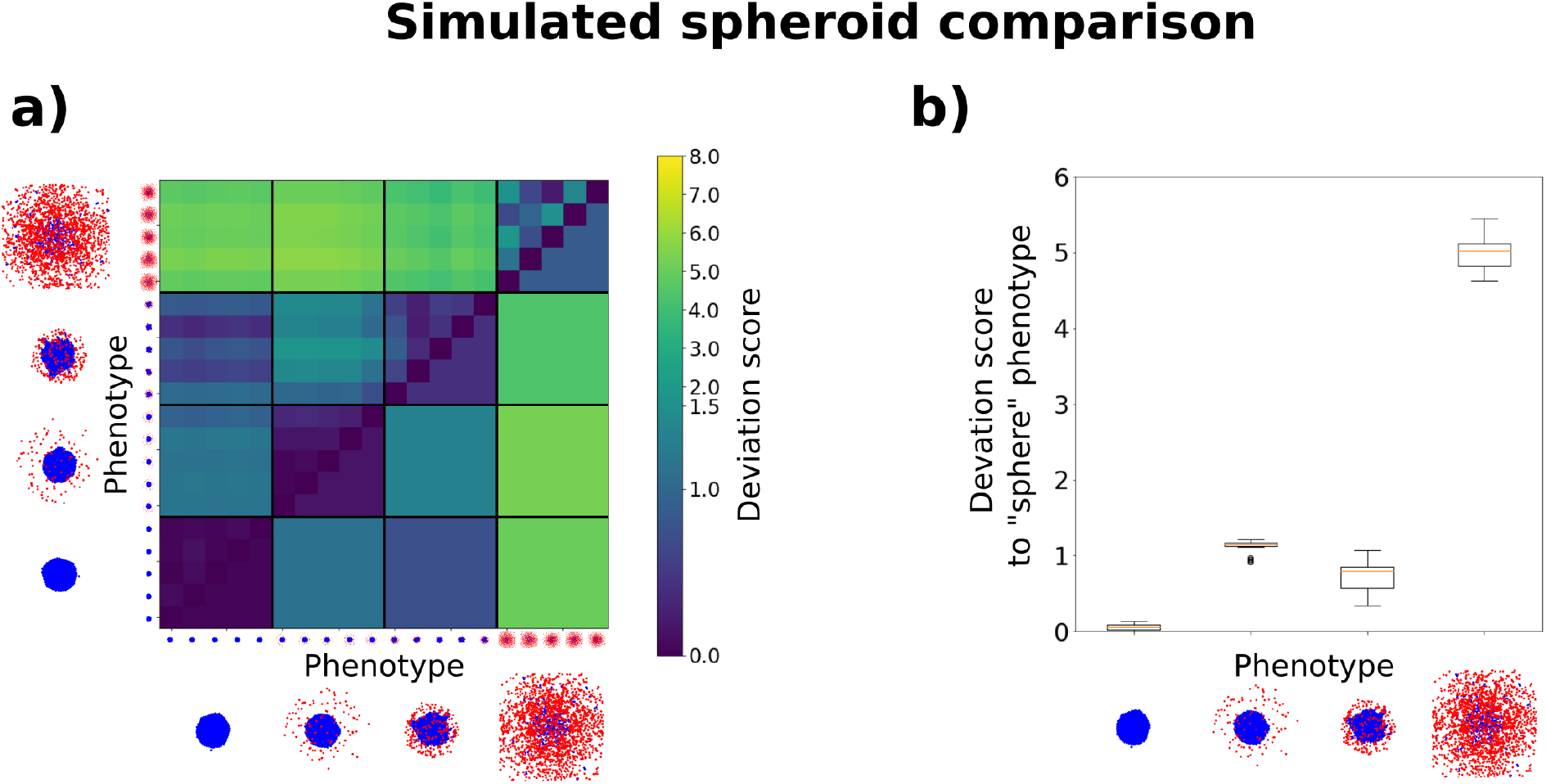
Deviation score comparison for four simulated spheroid phenotypes. **a)** Shown are the deviation scores for five replicates of each phenotype on the upper triangle, and the average deviation score over all replicates of each phenotype on the lower triangle. We observe the highest deviation between the “disordered” phenotype and the others, with the maximum deviation between the “spherical” and the “disordered” phenotypes. The “spherical”, “spherical with far gaslikes” and “deformed” phenotypes, which are more similar from a visual perspective, show a smaller deviation score using our analysis, but are nonetheless distinguishable. For illustrative purposes we have stretched the color bar for deviation score values below 1.5. **b)** Box plots of the deviation score values between the “spherical” phenotype and each other phenotype. The values used here correspond to those used for the lowest row of subfigure a). We observe that the deviation scores for the “spherical” phenotype compared with the other phenotypes consistently lie above the maximum deviation score of the “spherical” phenotype compared with itself.

### 2.6 Validation: comparing experimental spheroids

To demonstrate the fact that our deviation score can also be used for experimental data, we again turned to the dataset on which we based our initial simulations. Kang et al. previously investigated the invasive behavior of tumor spheroids cultured in graded collagen concentrations [13]. Their data set contained cell-resolved 3D snapshots of spheroids in four different collagen concentrations (1-2-3-4 mg/ml) at different times (days 0-1-2-3) during invasion. For each collagen concentration and day of culture, data from three individual spheroids were acquired using a combination of optical clearing and multiphoton microscopy. Since the optical clearing procedure requires fixation, data from successive days of culture share the same initial conditions but do not originate from the same spheroid (for more details, see Section 4.4).

In order to keep the scale consistent throughout this study, we used the same standardization and weight factors as for our simulated spheroids. We compared the spheroids grown for one day, two days, and three days, and visualized them in Fig 7. Similar to Fig 6, in subfigure a) we show, for each growth duration, the deviation scores for individual replicates on the upper triangle, and the average deviation score over a set of replicates on the lower triangle. For illustrative purposes and consistency, we have again stretched the color bar for deviation score values below 1.5. In subfigure b), we show box plots of the deviation score values between collagen 1 and each other concentration. As expected, we observe relatively low devation between the day 1 spheroids, both in subfigure a) and b). This changes at day 2, where the difference between collagen concentrations becomes clearer. However, while the difference is visible in the average deviation score, there is still strong overlap between the values across concentrations, as seen in subfigure b). Finally, on day 3 we see the highest difference between collagen 2 versus the other concentrations, while spheroids of collagen density 3 and 4 are most similar to each other. This is consistent with qualitative observations from Fig 2 a) and from prior quantifications by Kang et al. (cf., Figure 4c in [13]) who observed a sudden transition in single cells individualization during invasion between collagen cncentrations of 2 and 3 mg/ml.

**Figure 7:**
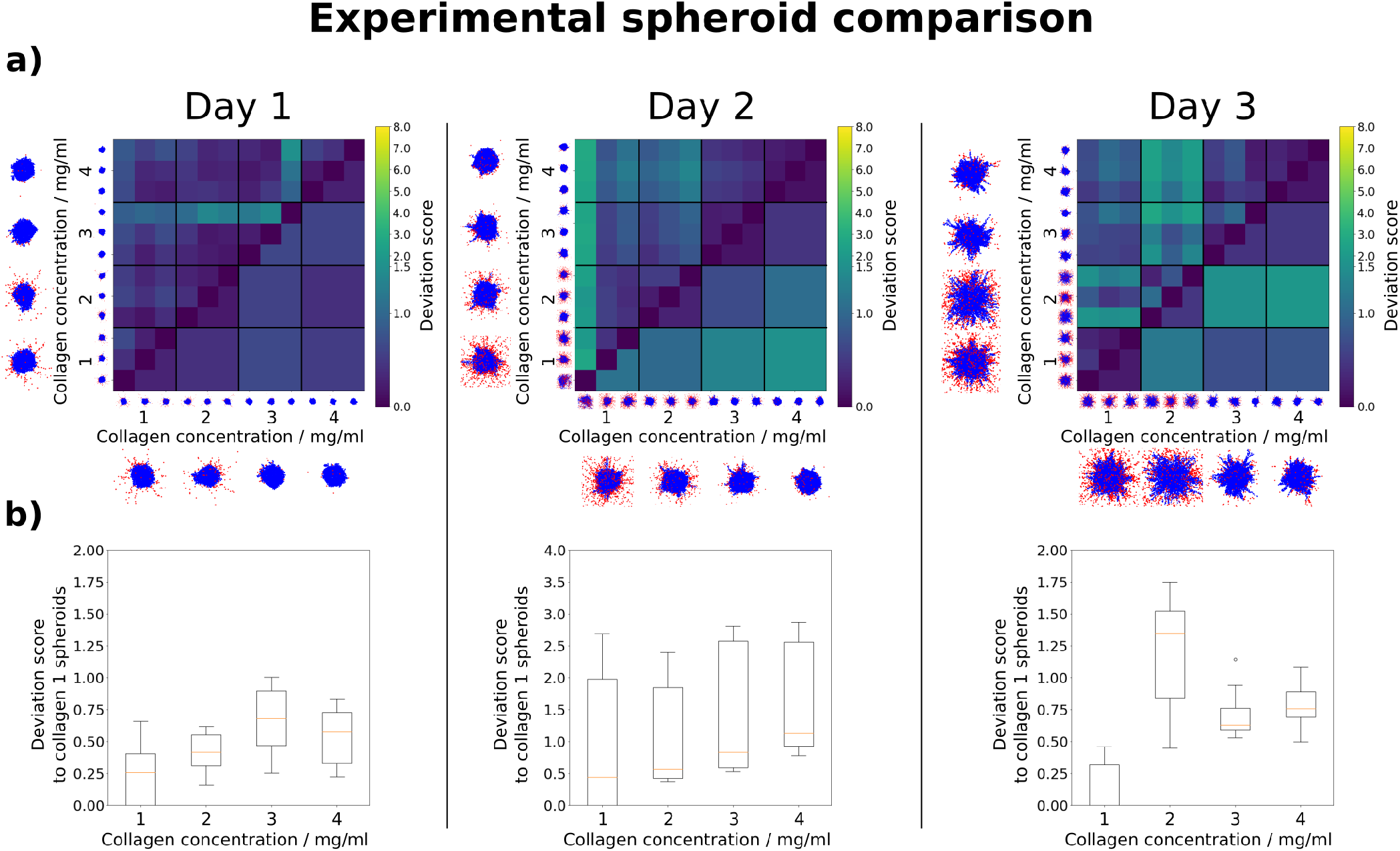
Deviation score comparison for *in vitro* spheroids cultured in four collagen concentrations *c* (data provided by Kang et al [13]). **a)** Comparison of spheroids invading for one, two and three days respectively. For each day, the deviation scores for three replicates of each collagen concentration are shown on the upper triangle, and the average deviation score for each collagen concentration is shown on the lower triangle. Due to matching initial conditions, we observe low deviation scores between spheroids grown for one day. These differences increase at day 2, where we observe an approximately linear increase of the average deviation score from *c* = 1 mg/ml to *c* = 4 mg/ml. Finally, at day 3, we observe the lowest deviation between *c* = 3 mg/ml and *c* = 4 mg/ml. This underlines the findings by Kang et al., who observed a transition in invasion behavior between 2 mg/ml and 3 mg/ml. **b)** Deviation score box plots from spheroids grown for one, two and three days respectively. The box plots for each day show the deviation score values between *c* = 1 mg/ml and each other concentration. These values correspond to those used for the lowest rows in a). We observe that for spheroids grown for one and two days, the deviation score values of *c* = 1 mg/ml compared with itself are similar to the deviation score values of *c* = 1 mg/ml compared with the other concentrations. Therefore the differences are not sufficient to clearly distinguish between them. This changes at day three, where each concentration shows a higher deviation score to *c* = 1 mg/ml than *c* = 1 mg/ml compared with itself.

### 2.7 Comparing simulated and experimental spheroids

Finally, as a precursor to our future work, we applied our analysis method to the comparison between simulated and experimental spheroids. For this, we used both the test cases from our simulated phenotypes (see Section 2.5), and the experimental data from spheroids grown for three days (see Section 2.6). Since the simulation parameters used here were not fitted to the data, but represented default parameter sets, we did not expect a high degree of similarity. On the other hand, this provided an opportunity to investigate both the overall deviation score and the underlying feature distances, and to demonstrate how the differences in spheroid morphology manifested themselves within the features. In subfigure a) of figure 8, we show the comparison between each experimental replicate (horizontal) and each simulated replicate (vertical) on the left side. On the right side we show the average within replicates. Here, we observed the highest deviation scores between collagen density 2 and “spherical with far gaslikes” (SFG) spheroids. Visually, this is sensible when comparing the 2D images of the replicates; the shape of the SFG spheroids differs strongly from that of spheroids in 2 mg/ml collagen, as does the number and location of cells classified as gaslike. Furthermore, we observed the lowest deviation scores between spheroids in 4 mg/ml collagen and “deformed” spheroids. Here, the spheroid shape visually matched much better between the replicate images. These visual differences and similarities are reflected in our features, as seen within subfigure b). Here we decomposed the overall deviation score back into its components, and thereby show the influence of each feature on it. We see that the distance between the collagen density 2 spheroids and the “spherical with far gaslikes” spheroids is noticable for all features except for the cell volume distribution. Of these, the gaslike cell distribution and the spheroid surface area exhibit especially high distances. On the other hand, collagen density 4 spheroids and “deformed” spheroids match comparatively well for each feature, and the spheroid surface deformation distance between them is the lowest of all.

**Figure 8:**
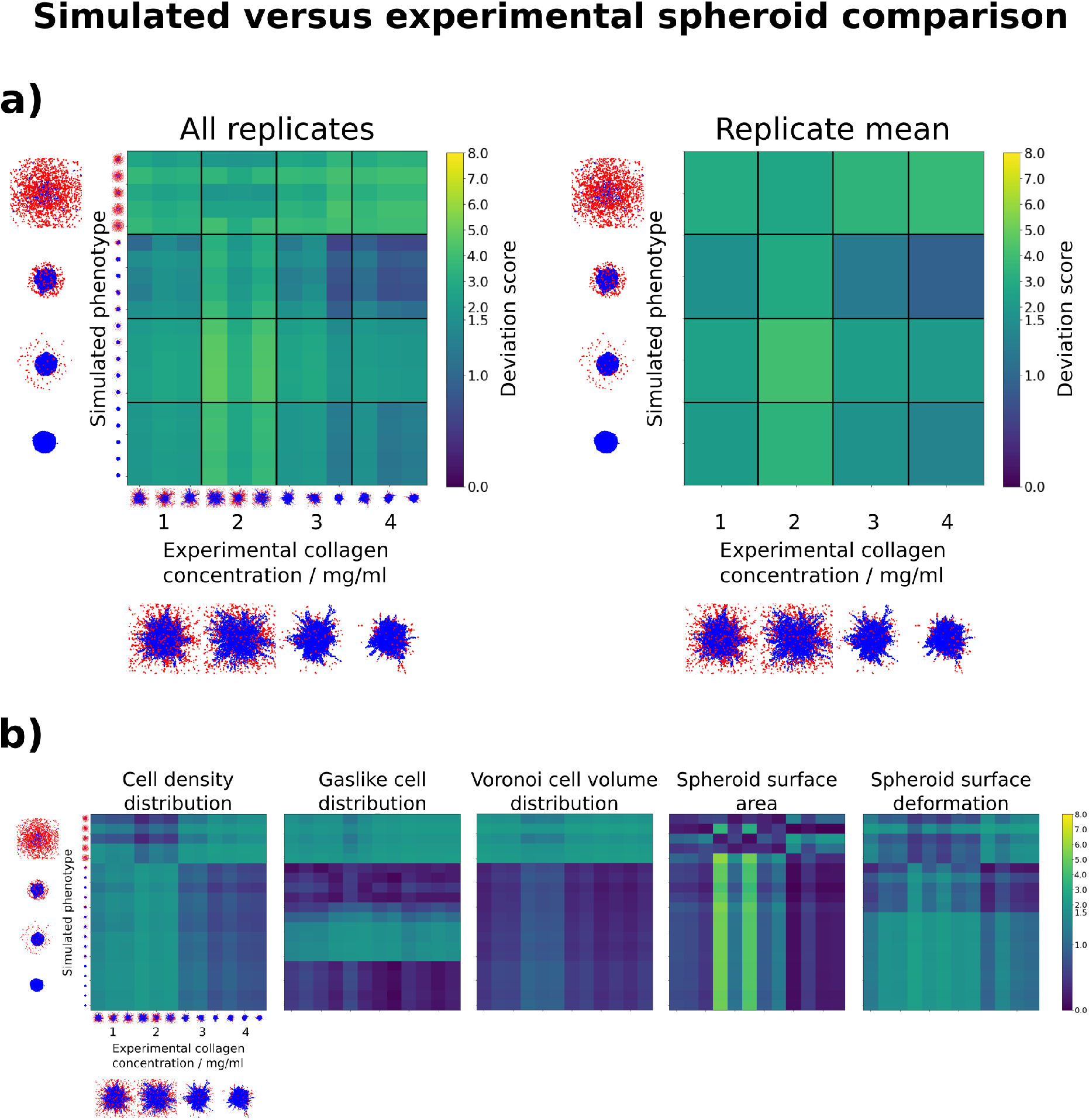
Deviation score comparison between *in vitro* spheroids grown in media at four different collagen concentrations *c* (data provided by Kang et al [13]), and *in silico* spheroids exhibiting four different phenotypes, simulated by us. **a)** Shown are the deviation scores between three replicates of each collagen concentration, grown for three days, and five replicates of each simulated phenotype, simulated for 250 000 MC steps. Each individual deviation score is shown on the left, and the average within a pairing of collagen concentration and phenotype is shown on the right. We observe the highest average deviation score between *c* = 2 mg/ml and the “spherical with far gaslikes” phenotype. The lowest average deviation score is found between *c* = 4 mg/ml and the “deformed” phenotype **b)** Individual metric distances for each of the features constituting the overall deviation score. Shown are the standardized metric distances between three replicates of each collagen concentration, grown for three days, and five replicates of each simulated phenotype, simulated for 250 000 MC steps. The highest deviation score observed in a) is a combination of high metric distances in all features, especially the spheroid surface and gaslike cell distribution. On the other hand, the lowest deviation score observed in a) stems from overall low values, especially in the spheroid surface deformation.

### 2.8 Nastjapy

During our derivation of the overall deviation score and its application to various data, we developed the Python package *nastjapy*. Through this, we wanted to facilitate the use of our procedure by others. The package can be found at http://www.gitlab.com/nastja/nastjapy. *Nastjapy* allows the investigation of spheroids and other single-cell resolved data from different origins. It thereby unifies the analysis pipeline for simulated data and data from multiple experimental sources. See Section 4.5 for more details.

## 3 Discussion and conclusions

Both experimentalists and theorists produce data concerning tumor spheroids. However, both the quantitative comparison between different experiments or simulations, and the process of fitting simulations to experimental data, are hindered by the lack of an adaptable distance measure that captures the similarity of the spatial features of two spheroids. We aimed to solve this issue via the following steps. First, we proposed a set of five relevant spatial features, which could be extracted from spheroid point clouds. Next, we devised metrics to compare each feature, and combined all metrics into an overall deviation score. We also provided an optimization scheme which could be used to adapt the deviation score to the specific use case. For this, we turned to four *in silico* spheroid phenotypes which emerged from our simulations, and used them to standardize and combine the metrics into the overall deviation score *D*_*i,j*_. We did this by weighing individual metrics differently while maximizing a phenotype separation property (see equation 7). We characterized the behavior of our features by applying four different transformations to a point cloud obtained from a spherical simulated spheroid. We were able to confirm rotational invariance by analyzing the “rotation” transformation, and monotony for the others. While the features related to the spheroid surface showed some instabilities for higher transformation strengths in the “noise” and “deformation” transformations, this only occured when the point cloud was so disordered that a solid core could no longer be defined. This represents a non-biological simulation scenario which lies outside of our domain of interest. Overall, the behavior of the features was therefore considerered suitable to quantitatively compare the structure of spheroids. Interestingly, the overall deviation score did not only scale monotonously with the strength of the studied transformations, but did so approximately linearly within the reasonable domains. This is a useful property for a distance measure, which was not used as a constraint, but instead emerged as a result of our optimization scheme.

During the investigation of our simulated spheroid phenotypes, we found that our deviation score distinguished well between dissimilar spheroid phenotypes, as all phenotypes showed the highest deviation score towards the “disordered” phenotype. This large distance was a value that was desired, as this phenotype is a non biological edge-case. Since we also observed non-negligible deviation between the other three phenotypes, we were able to confirm that our strategy is also applicable to more similar spheroids. We therefore consider *D*_*i,j*_ to represent a useful metric for the systematic quantification of spheroid similarity. This was further confirmed by our analysis of experimentally measured spheroids generated by Kang et al. [13].

One of our main goals in defining the deviation score was to have a reliable objective function for fitting simulated spheroids to experimental data. Therefore, as an outlook for our future work we included a comparison between simulations with unfitted, default parameters, and the aforementioned experimental spheroid data. Here, we were able to highlight, which features contributed most towards each deviation score, and to show that the quantities matched well with a visual comparison.

The features we have defined here admitedly have some limitations. They cannot, for example, measure the dynamics of cell movement over time. Furthermore, spheroids or other tissues we might want to apply this method to, may be composed of multiple different types of cells, and we currently do not distinguish between these. However, our strategy is meant to serve as a base to build upon, and since we implemented it in our freely available *nastjapy* framework (see Section 4.5), it can easily be extended. We aim to further develop this in the future, via incorporating more features. Points of interest would be generating features spanning multiple timesteps, e.g. cell velocity correlation and autocorrelation. Furthermore, we envision features which will enable the application of the analysis scheme to non-spheroid tissue models as well, e.g. cell composition analysis, cell cluster analysis, etc.

Our ultimate aim in working with *Cells in Silico* is building a cellular digital twin of a macroscopic tumor. The feature analysis which we developed here represents a promising step towards reaching this goal, because we will use it to fit parameters of CiS related to cell-cell adhesion, motility and ECM invasion, which will be done by iteratively comparing simulated and experimental spheroids. Thereby, we want to restrict these parameters before moving towards the simulation of larger tissues.

## 4 Methods

### 4.1 Model description

*Cells in Silico* is a framework for simulating the dynamics of cells and tissues at subcellular resolution, which was previously developed by our group [19]. It combines a Cellular Potts Model (CPM) at the microscale with nutrient and signal exchange at the mesoscale and an agent-based layer at the macroscale. This enables detailed capture of individual cell dynamics. Furthermore, as an extension of the *NAStJA* framework [25] its efficiency scales excellently with increasing system size and CPU core number. Hence, CiS has already been used for simulating tissues composing millions of cells [19]. Here, we briefly outline the main properties of the microscale, mesoscale and macroscale layers, and a more detailed description can be found in [19].

#### Microscale

The CPM was developed by Graner and Glazier in 1992 [26], as an extension of the Potts model. In it, a system of lattice points on a regular grid is propagated according to its overall energy. Cells are defined as aggregates of points of the same type, and the overall energy of the system is built of multiple components *E*_*i*_, which dictate the morphology of and interaction between the cells. Weighted by coupling factors *λ*_*i*_, they are combined into the following Hamiltonian:

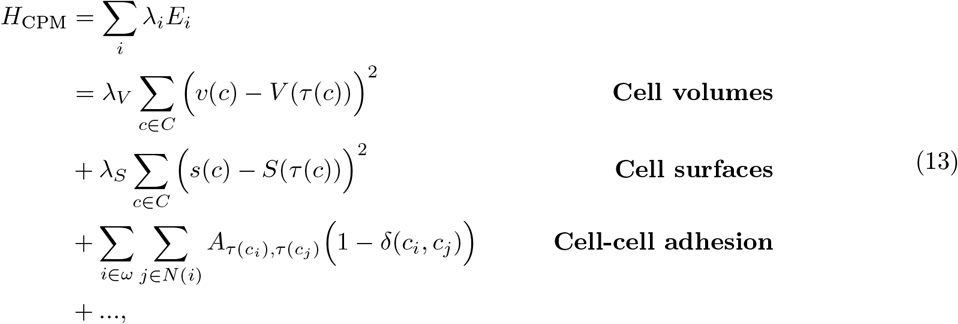

where *c* is a cell from the set of all cells *C, τ* (*c*) is the type of cell *c, s*(*c*) and *v*(*c*) are the current surface and volume of cell *c, S*(*τ*) and *V* (*τ*) are the target surface and volumes of cells of type *τ, A* is the adhesion coefficient matrix for all cell types, *N* (*i*) are all lattice points neighboring point *i*, and *δ* is the Kronecker delta. Equation 13 can be extended to include further effects, such as cell motility [27] (see also Section 4.2.

#### Mesoscale

CiS includes the capability of introducing signals or nutrients to the system. These can be exchanged between cells via the cell-cell interface. As this functionality is outside of the scope of this study, we only briefly mention it here.

#### Macroscale

While using the CPM layer allows for excellent reproduction of cell shape and deformation, there are other important cellular functions which are not intrinsically captured. For example, the CPM Hamiltonian does not in itself include the effect of cell division. Furthermore, while self-propelled cell motility can be added to equation 13 [27], the direction of the motility vector must be periodically updated for each cell, to ensure realistic movement, e.g. via random walk (see also Section 4.2). This requires information on the cell center location, which must be extracted from the CPM. A third aspect, which is very important for simulating realistic tumors, is the capability of *in silico* cell mutation. Here, cell parameters such as division rate, motility magnitude, cell-cell adhesion etc. must be adjusted at the time of division. All the aforementioned aspects are treated in the macroscale layer. It combines information gathered from the lower layers with higher-level parameters, which results in an agent-based system. Here, the conditions for cell division are checked, the division process is carried out, the motility direction is updated, etc.

By combining micro-, meso- and macroscale, we gain a versatile tool, which can then be parameterized.

### 4.2 Model parameters

Using CiS, we simulated a multitude of spheroids. We based our simulations on experimental spheroid data provided by our collaborators [13]. Hence, each simulated spheroid had an initial diameter of 200 µm, contained roughly 2000 cells, and was placed in the center of a volume spanning 800 × 800 × 800 µm^3^. Using CiS, we propagated this system at a range of different simulation parameter combinations, which are highlighted below.

#### Extracellular matrix

The extracellular matrix (ECM) is a scaffold within tissues, which connects cells and serves both as a structural component and cell maintenance network [28]. It is composed of overlapping fibrous polymers, such as collagens, proteoglycans and glycoproteins [29]. Tumor spheroids are often placed into a collagen matrix, which serves as a proxy for an *in vivo* ECM [13, 30]. To capture this in our simulations, we modeled the ECM as overlapping, randomly oriented fibers, which were placed within the system volume surrounding the spheroid (see supplementary Fig 1). These fibers represented rigid obstacles for the cells, to which they could adhere, but which could not be displaced. Since in reality the ECM is not a solid structure, but can be modified and degraded by cells [31], this alone was an overly simplified description. We therefore added a degradation effect, by which cells removed ECM lattice points with which they were in direct contact on the CPM lattice. This occured after a set number of MC steps, as described by the ECM degradation period parameter. During each degradation event, a lattice point in contact with a cell was removed with a probability of 50 %. For our simulations, we varied both the ECM density and the ECM degradation period. The ECM density was varied between volume fractions of 0 % and 90 %. The ECM degradation was either disabled, or its period was varied between 1 000 and 10 000 MC steps.

#### Cell-cell adhesion

Changes in the adhesion strength between cells are a well known factor which facilitates invasion [7]. We therefore varied the adhesion strength parameter within our simulations, by changing the adhesion coefficient matrix in the third component of equation 13. This matrix describes the strength of adhesion between different cell types, as well as the strength of adhesion between cells and the ECM.

#### Cell motility magnitude and persistence

Similar to cell-cell adhesion, the cell motility is strongly connected to the invasion properties of cells [32]. To include it, we modified the CPM Hamiltonian by adding a directional potential to each cell with the following contribution [27]:

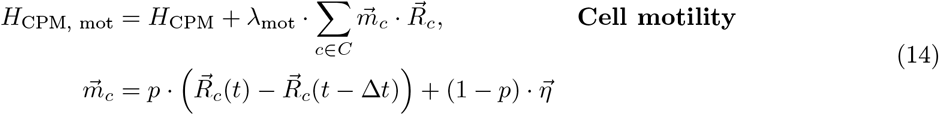

where 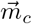 is the potential and direction a cell *c* experiences, *R*_c_ is the center of mass of cell *c*, and 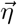 is a random vector obtained by a Wiener process [33]. The motility is implemented as a modified persistent random walk of each cell. The energy contribution of each cell is the dot product of 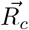 and 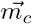, which in turn is determined by the cell’s previous movement and 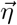. The mixture of persistent and random movement can be chosen by the persistence parameter *p* ∈ [0, 1]. Cells with *p* = 0 perform purely random walks, and cells with *p* = 1 perform purely persistent walks. The coupling strength of this energy term to the CPM is given by *λ*_mot_.

### 4.3 Emerging phenotypes

We observed four spheroid phenotypes which emerged from various parameter combinations. These were the following:

#### Spherical

Cell-cell adhesion dominated, and the cells remained in a spherical arrangement, with a relatively smooth surface of the spheroid bulk throughout the simulation.

#### Spherical with far gaslikes

Due to a low value of the motility magnitude parameter, cells did not move far on their own. However, through a combination of high ECM degradation rate and cell-ECM adhesion, cells in the outermost layer of the spheroid were pulled into the ECM. This occured at high ECM densities. The spheroid bulk remained intact.

#### Deformed

The cell-ECM and cell-cell adhesion parameters were at similar strength. Therefore, the spheroid retained a bulk structure, but lost its spherical shape at high ECM-density.

#### Disordered

The cells dissociated from each other due to high self-propelled motility, low cell-cell adhesion and low ECM density. Subsequently, the spheroid integrity was lost.

### 4.4 Experimental spheroid preparation and analysis

All experimental methods were previously reported by Kang et al. [13] and are briefly summarized here. Multicellular tumor spheroids were formed by seeding highly invasive, triple-negative MDA-MB-231 breast cancer cells [34] in low-attachment 96 well plates (Corning, No. 07201680) in the presence of 2.5% v/v Matrigel (Corning, No. 354234) [35]. Using this approach, ∼1000 cancer cells coalesced into a spherical aggregate (i.e., a tumor spheroid) of 300-to-400*µ*m in diameter over the course of 48 hours. Once formed, individual spheroids were fully embedded into a 3D fibrous gel prepared using rat-tail collagen I (Corning, No. 354249) [36]. As shown in Kang et al. [13], by varying the collagen concentration between 1 and 4 mg/ml, one can tune the fiber density and overall mechanical properties of the collagen network surrounding each tumor spheroids. MDA-MB-231 spheroids were then cultured in such 3D micro-environments for either 1 hour (Day 0), 24 hours (Day 1), 48 hours (Day 2), or 72 hours (Day 3). While at Day 0 all cells remained within the main spheroid (solid-like phase), over the course of 3 days tumor spheroids progressively developed strikingly different patterns of invasion as a function of collagen concentration, including single cell invasion in 1-2 mg/ml collagen (gas-like phase) and collective invasion in 3-4 mg/ml collagen (liquid-like invasion) [13]. For each time point, spheroids were fixed, optically cleared [37], stained with DAPI (Fisher Scientific, No. D1306), and imaged using a Bruker Ultima Investigator multiphoton microscope equipped with a long working distance 16x water-immersion objective (Nikon, 0.8 N.A., 3mm working distance) to enable whole-spheroid imaging [13]. The 3D positions of DAPI-stained cell nuclei were finally identified using a custom Matlab code developed by Kang et al. [13] and used herein as point cloud data to extract features from experimental spheroids. In this work we used point cloud data from spheroids imaged at days 0-1-2-3 in collagen concentrations of 1-2-3-4 mg/ml, *n* = 3 per group, except for day 0 in 1 mg/ml (*n* = 2), and day 2 in 2 mg/ml (n = 9).

### 4.5 Nastjapy

To facilitate the use of the analysis pipeline presented in this study, we developed the Python package *nastjapy*, which can be found at http://www.gitlab.com/nastja/nastjapy. For this study, *nastjapy* served three functions:

1. providing a unified interface for processing data from multiple different sources
2. performing efficient and parallel feature extraction and analysis
3. adaptating and computing the deviation score for specific applications.

*Nastjapy* allows treating spheroids and other single-cell resolved data from different sources, thereby unifying the analysis pipeline for simulated data and data from multiple experimental sources. Data from different experiments may come in different file types, i.e. CSV, HDF5, SQLite and matlab. Even files of the same filetype do not necessarily have to share their formatting between different experiments. *Nastjapy* provides a common interface between those sources and is written to be easily extendable for new file types or formats. Extracting all features for many different spheroids, possibly at multiple points in time, as well as the comparison between a large number of spheroids, can quickly become computationally expensive. *Nastjapy* therefore provides efficient implementations of all compute-intense functions, as well as the option to parallelize the computation on multiple cores via MPI. Lastly, *Nastjapy* offers all functions neccessary to use the methods described in this study to adapt and compute the the deviation score for a given application. Some further analysis tools and visualization capabilities area also provided.

## 5 Acknowledgement

The authors gratefully acknowledge the Gauss Centre for Supercomputing e.V. (www.gauss-centre.eu) for funding this project by providing computing time through the John von Neumann Institute for Computing (NIC) on the GCS Supercomputer JUWELS at Jülich Supercomputing Centre (JSC).

The present contribution is supported by the Helmholtz Association under the joint research school “HIDSS4Health – Helmholtz Information and Data Science School for Health.

---

**Supplementary Figure 1:**
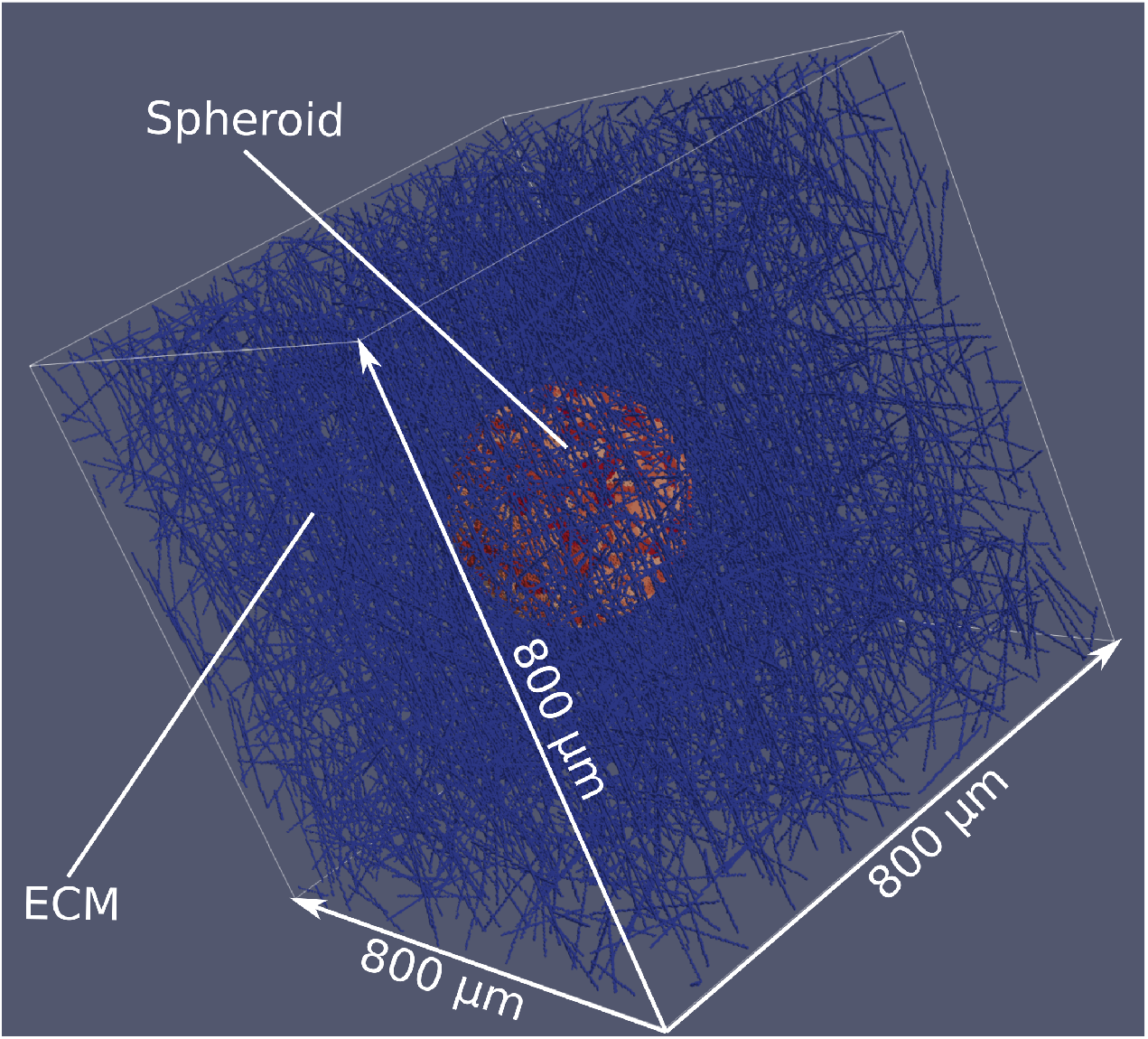
Visualization of initial conditions of spheroid simulations. Each spheroid was placed in the center of an 800 × 800 × 800 µm^3^. volume, and surrounded by an ECM of varying density (see Section 4.2). For illustrative purposes, the ECM density was set to a low value here, such that the spheroid is still visible.

**Supplementary Figure 2:**
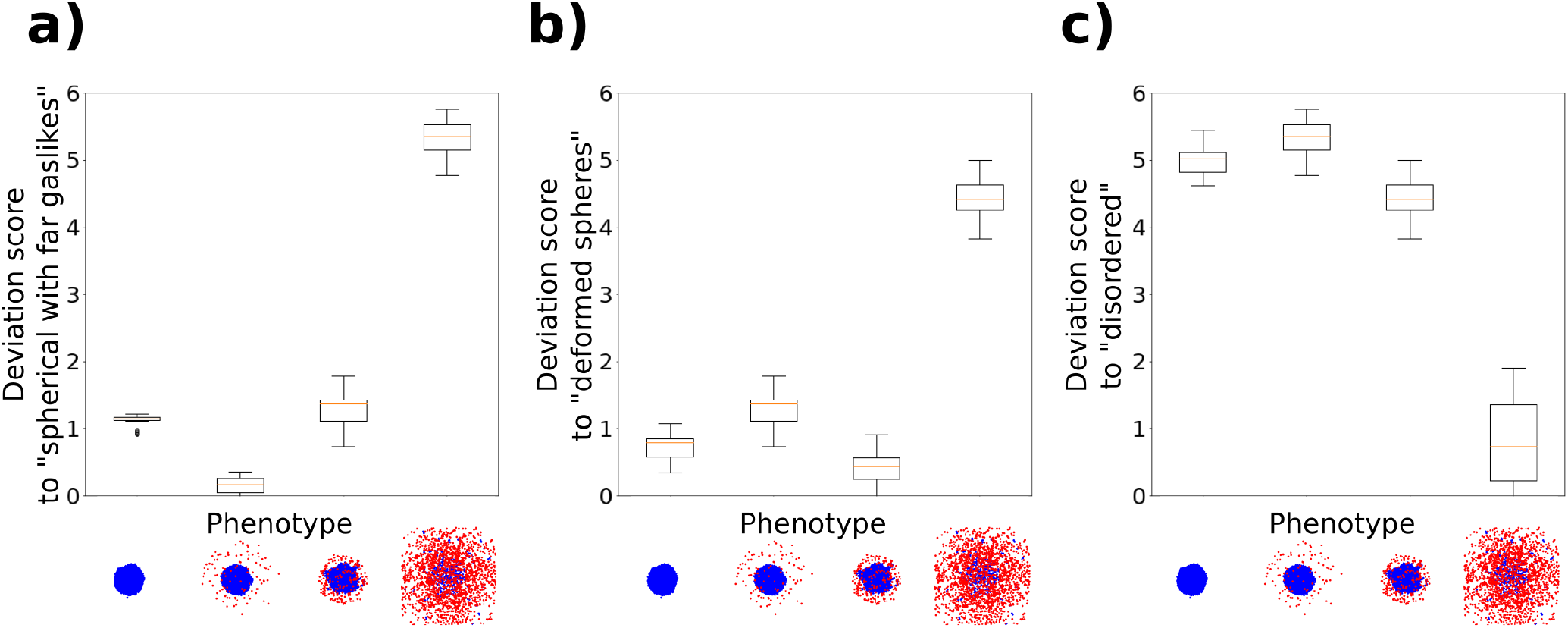
Deviation score box plots for simulated phenotypes.**a)** Each phenotype compared to the “spherical with far gaslikes” phenotype. **b)** Each phenotype compared to the “deformed” phenotype. **c)** Each phenotype compared to the “disordered” phenotype.

**Supplementary Figure 3:**
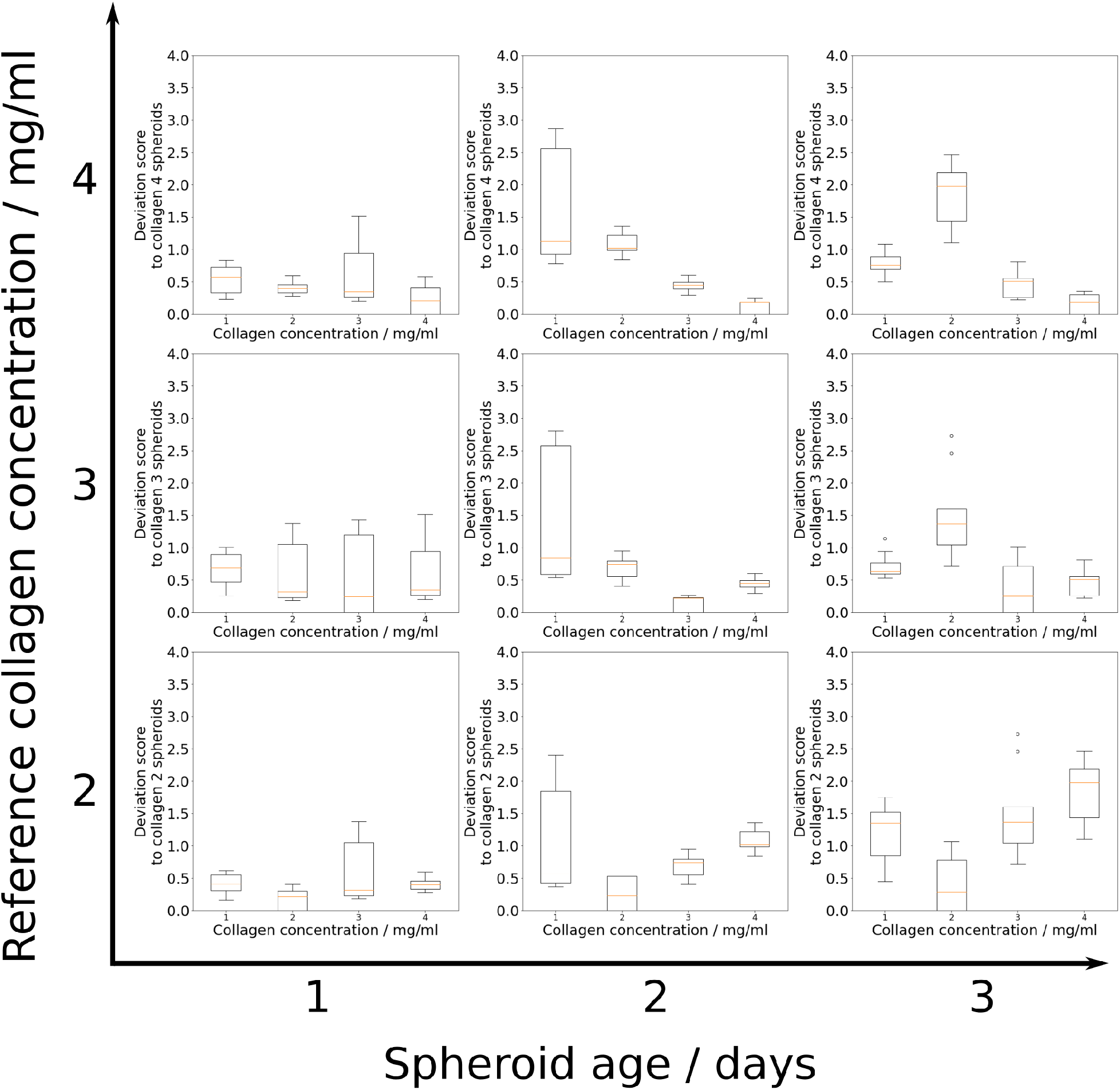
Deviation score box plots of *in vitro* spheroids grown in media at four different collagen concentrations (data provided by Kang et al [13]).

## Notes

### Competing Interest Statement

The authors have declared no competing interest.

